# A broad-host-range event detector: expanding and quantifying performance across bacterial species

**DOI:** 10.1101/369967

**Authors:** Nymul Khan, Enoch Yeung, Yuliya Farris, Sarah J. Fansler, Hans C. Bernstein

## Abstract

Modern microbial biodesign relies on the principle that well-characterized genetic parts can be reused and reconfigured for different functions. However, this paradigm has only been successful in a limited set of hosts, mostly comprised from common lab strains of *Escherichia coli*. It is clear that new applications – such as chemical sensing and event logging in complex environments – will benefit from new host chassis. This study quantitatively compared how a chemical event logger performed across multiple microbial species. An integrase-based sensor and memory device was operated by two representative soil Pseudomonads – *Pseudomonas fluorescens* SBW25 and *Pseudomonas putida* DSM 291. Quantitative comparisons were made between these two non-traditional hosts and two bench-mark *Escherichia coli* chassis including the probiotic Nissle 1917 and common cloning strain DH5α. The performance of sensor and memory components changed according to each host, such that a clear chassis effect was observed and quantified. These results were obtained via fluorescence from reporter proteins that were transcriptionally fused to the integrase and down-stream recombinant region and via data-driven kinetic models. The *Pseudomonads* proved to be acceptable chassis for the operation of this event logger, which outperformed the common *E. coli* DH5α in many ways. This study advances an emerging frontier in synthetic biology that aims to build broad-host-range devices and understand the context by which different species can execute programmable genetic operations.

## INTRODUCTION

Synthetic biology is built on the concept that complex biological behaviors can be programmed using relatively simple modules of biological parts. While the field of microbial biodesign has seen major advances, the overwhelming majority of parts have only been tested in model organisms. To date, we know little about how even our most standard genetic devices will perform in microbial hosts beyond common laboratory strains of *Escherichia coli* or *Saccharomyces cerevisiae*. This represents a major knowledge gap and limitation in the field. While useful for the development and demonstration of capabilities under stable laboratory conditions, these species do not survive well in many real-world applications. Most traditional microbial hosts have limited metabolic potential, preferring substrates such as simple sugars that are typically not available in environments relevant to the next generation of synthetic biology applications such as event detection within soils, built environments or the human gut. Therefore, programmable genetic devices must be expanded into new, non-traditional chassis that are already evolved to operate in complex, dynamic environments.

One of the most common biodesign principles is that well-characterized genetic parts – e.g., promoters, UTRs and transcription factors – can be reused and reconfigured to program different functions. Some of the benchmark examples are given by the toggle switch (1), repressilator (2) and previous demonstrations of integrase-based recording devices (3–6); all of which were exclusively demonstrated in *E. coli.* These devices have laid the foundation for more applied microbial sensor-regulator-actuator devices that have been developed to detect/report signals from the mammalian gut (7,8) and chemical threats (9,10); yet even these advanced examples relied solely on the genetic tractability of *E. coli*. Synthetic biologists are keen to harness new non-traditional hosts such as *Pseudomonads* (11–13); yet, successful transplantation of broad-host-range genetic devices across multiple bacterial species has remained elusive, until now.

Here we present a study that demonstrates how a relatively simple chemical event logger performs across multiple microbial hosts. We chose to comparatively quantify each component of an integrase-based sensor/memory device between two *Pseudomonas* species – *Pseudomonas fluorescens* SBW25 (*Pf*) and *Pseudomonas putida* DSM 291(*Pp*) – along with two more standard *Escherichia coli* strains including the probiotic Nissle 1917 (*EcN*) and common cloning strain DH5α (*Ec*). The event detector was expressed from each host as the same sequence on an identical broad-host range expression vector. Here, we show that a genetically identical chemical event logging device can be ported across multiple species, including two Pseudomonads that open new chemical sensing/logging applications soil and plant-associated environments. The performance for each component of the device depended on the host – it was subject to a strong chassis effect. Hence, study presents a new broad-host range event logging system, which advances to a rapidly growing frontier in synthetic biology aimed at engineering devices that can function across multiple species and environments (14,15).

## MATERIALS AND METHODS

### Bacterial strains and cultivation

The bacterial strains used in this study includes *Escherichia coli* DH5α (New England Biolabs), *Escherichia coli* Nissle 1917 (isolated from probiotic Mutaflor capsule), *Pseudomonas fluorescens* SBW25 and *Pseudomonas putida* DSM 291 (DSMZ). All bacteria were cultured in Lauria-Bertani medium at 30 °C.

### Transformation

*E. coli DH5α* was transformed using standard chemical transformation protocol. For the other species, electrocompetent cells were prepared as follows: overnight cultures were diluted 1:100 into 200 mL LB medium and grown to OD600 nm of about 0.3-0.4 (mid-log phase); cultures were harvested and spun down in four 50 mL centrifuge tubes at 5000 × g and the supernatant was discarded; cell pellets were resuspended in 15% glycerol, combined into one 50 mL centrifuge tube and collected via centrifugation at 5000 × g. This wash cycle was repeated twice and the final cell pellet was resuspended in 1 mL 15% glycerol for electroporation. The cells were then transformed by electroporation at 12500 V/cm (200 Ω and 25 µF) in 1 mm cuvettes. The entire protocol was successfully carried out at room temperature as outlined in the Tu et al. 2016 study (16). The efficiency, in general, was found to be higher in the room-temperature methods than in conventional ice-cold methods.

### Plate reader and cytometry assays

For each bacterial species, three positive transformants were grown, passaged twice at 30 °C and then assayed in a 24-well plate at eight different IPTG concentrations (0, 0.01, 0.05, 0.1, 0.2, 0.5, 0.8 and 1 mM). Each well of the plate contained 1.8 mL LB + kanamycin (50 µg/mL) + IPTG. A Synergy H1 (Biotek, Winooski, VT) was used as the fluorescent plate reader for all assays. 1 µL samples were collected from each of the wells at various stages of growth and analyzed via flow cytometry (Novocyte, ACEA biosciences, San Diego, CA). Simultaneously, 100 µL samples were also collected and frozen at −80 °C. Plasmid DNA was later extracted from the frozen samples and used for real time qPCR assays to measure the fraction of device flipped.

### Real time quantitative PCR

Real time PCR was performed by ARQ Genetics LLC (Bastrop, TX) on the BioRad CFX384 Real Time System (BioRad, Hercules, CA) using assays specific for each plasmid. All of the plasmid DNA was extracted with a Zyppy – 96 Plasmid Miniprep kit (Zymo, Irvine, CA) following the next steps: all of the stains were grown at 30 °C and harvested at 3-5 h intervals; the DNA was quantified by performing PicoGreen assay on the Biotek Synergy H1 (Biotek, Winooski, VT) and reactions were diluted to matching concentrations. Each reaction within multi-plate wells contained 5 μL of TaqMan Universal Master Mix II (Applied Biosystems, Waltham, MA), 2 μL of each sample template and 0.5 μL of each specific plasmid assay in a reaction volume of 10 μL. Cycling conditions were as follows: 95 °C for 10 min for polymerase activation, followed by 40 cycles of 95 °C for 15 seconds and 63 °C for 1 min. Data analysis was performed using CFX Manager software from BioRad, version 3.1. The experimental Cq (cycle quantification) was calibrated against the standard curve for each plasmid orientation.

### Numerical simulation

The system of ODEs were solved numerically using the ‘deSolve’ package (17) in R (18). The results were fitted to experimental data to estimate the five kinetic rate constants (*P*_*A*_, *P*_*B*_, *P*_*C*_, *D* and *k*_*flip*_) for each of the species in the model. The specific growth rate (*µ*) was calculated from measured OD600 nm at each time point and given as an input to the numerical solver.

## RESULTS

### The broad-host range device and its components

A two-state chemical event logger was built and quantitatively compared across microbial hosts to determine the chassis effect on performance. The device was built with specific sensor and memory components (Fig. 1a). The sensor apparatus consisted of an IPTG inducible *P*_*lac*_ promoter (with *lac* operator) driving the expression of the Bxb1 serine-integrase. The Bxb1 gene was transcriptionally fused to a green fluorescent protein (GFP) to monitor the sensor’s output. The *lacI*^*q*^ transcription factor, controlling the induction of *P*^*lac*^, was driven by the constitutive promoter, 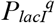. The memory element had two potential states that depended on the Bxb1 integrase. The “*off state*” was the initial or unchanged state of the plasmid where the constitutive *P*_*tac*_ promoter (without a *lac* operator) was in the reverse orientation from its intended open reading frame, followed by a unique barcode DNA sequence. This construct was enclosed by the *attB* and an *attP* recombination sites recognized by Bxb1. The induced state or “*on state*” was controlled by the formation of a mature Bxb1 dimer, DNA binding and tetramer formation (19), and the respective recombination of *attB* and *attP*. This process re-oriented the constitutive *P*_*tac*_ to drive expression of a red fluorescent protein (RFP). A permanent digital memory output was stored by the orientation of the barcode. The performance of the sensor and logger elements were measured by the respective GFP and RFP signals. The entire device was built into a single contig and cloned into the broad-host-range vector pBBR1MCS2 (20) (Fig. 1b).

**Figure 1.**
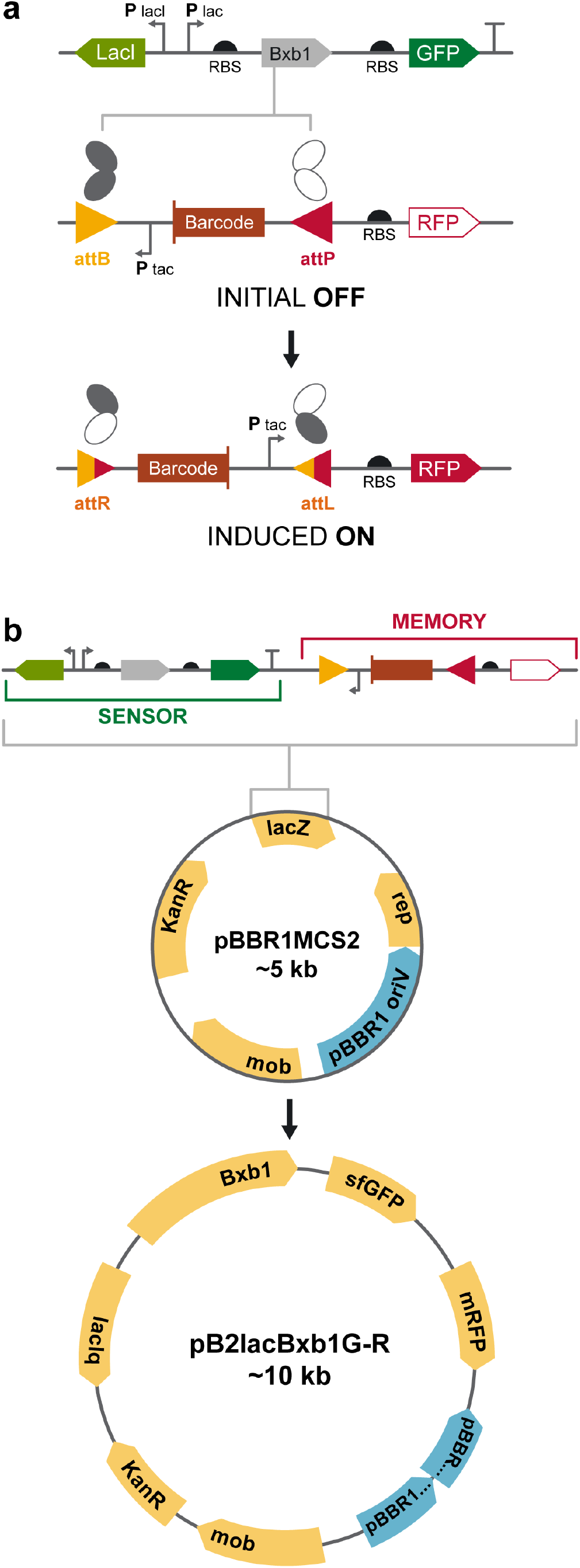
The broad-host range event logger and its modes of operation. (a) Expression of the Bxb1 integrase was controlled by the Plac promoter and respective IPTG concentrations. Mature Bxb1 proteins dimerize and bind to DNA recombination sites (attB and attP). The dimers on the two ends of the recording element join to form a tetramer in a synaptic event that folds the DNA in the process. DNA strands are exchanged when Bxb1 monomers trade positions, flipping the internal region, which contains both a barcoded digital recorder and a constitutive Ptac promotor. The new sites, attL and attR-formed in the process of DNA flipping - can no longer stay attached to the Bxb1 dimers because of altered sequence, therefore release the dimers in an irreversible digital recording process. GFP and RFP reporter genes were transcriptionally fused onto the IPTG inducible sensor and recording recombination sites. (b) Map of the device with the sensor and memory components cloned in place of the lacZ gene on the pBBR1MCS2 broad-host-range vector.

### Quantifying the chassis effect

The device was operational across each of the four species. Performance of each component – sensor and logger – was assayed at eight different IPTG concentrations (0 – 1 mM) by measuring the respective mean GFP and RFP signals (Fig. 2). Total growth was also measured simultaneously via optical density (OD600 nm). The specific growth rates (μ) showed that each species – except *EcN* (μ = 1.087 ± 0.017 h^−1^) *–* had very similar growth rates under these conditions: *Ec* (0.427 ± 0.016 h^−1^), *Pf* (0.508 ± 0.019 h^−1^) and *Pp* (0.439 ± 0.0436 h^−1^) (Supplemental Fig. S1a). The standard deviations represent the variation in growth rates across all IPTG treatments and ranged from approximately 1.5% to 10% of mean values. The induction strength of the device – controlled by IPTG concentration – showed minimal effect on the specific growth rate. While this indicates that there was little additional metabolic load with respect to IPTG induction, the plasmid encoded device itself imposed a significant metabolic burden on both *Pseudomonas* hosts. This was apparent from the wild-type growth rates of these species, measured at 0.759 ± 0.017 h^−1^ and 1.127 ± 0.012 h^−1^ for *Pf* and *Pp* respectively (Supplemental Fig. S1b); much higher than the respective engineered strains. In contrast, there was very little change in specific growth rate of the engineered *E. coli* hosts compared to their respective wild-types (0.412 ± 0.012 h^−1^ and 0.979 ± 0.022 h^−1^ for *Ec* and *EcN* respectively).

**Figure 2.**
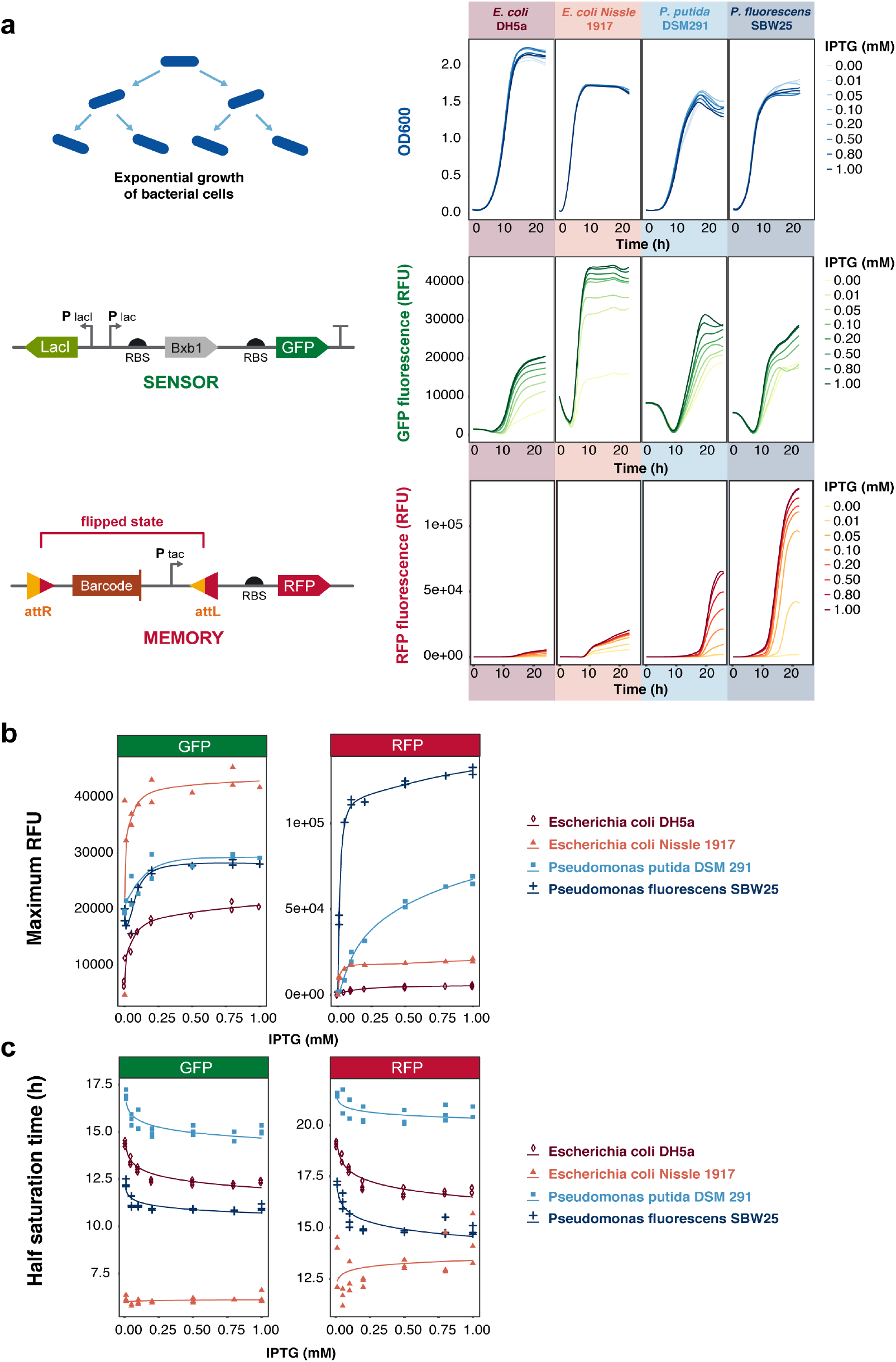
Quantifying the chassis effect. (a) The top four panels represent cell growth (OD600 nm), at different IPTG concentrations, of E. coli DH5α, E. coli Nissle 1917, P. putida DSM 291 and P. fluorescens SBW25 from left to right respectively. The middle row of graphs represents the fluorescence output of the GFP reporter that was transcriptionally fused to the sensor element; the bottom row represents the fluorescence output from the RFP that was fused to the memory component. (b) The maximum GFP and RFP fluorescence at different IPTG concentrations. (c) The activation coefficient shown as half-saturation times of GFP and RFP at different IPTG concentration (> 0 mM IPTG).

The sensor apparatus did not show the tight transcriptional control that was expected from the *P*_*lac*_/*lacI* system in any of the hosts (Fig. 2a). All species had significant level of basal GFP fluorescence (at 0 mM IPTG), indicating leaky expression. Among all the hosts, the sensor component performed the best in *EcN* which had high fluorescence and low basal expression, indicating higher response and tighter regulation than others. *Pf* and *Pp* had similar but high basal GFP expression. While natural fluorescence of the Pseudomonads in the GFP emission spectrum was initially thought to influence the reporter signals, detailed examination revealed that wildtype fluorescence was insignificant compared to the devices’ GFP signal. *Ec* had the lowest GFP fluorescence of all the species but also the lowest basal expression. In general, however, the *E. coli* species showed tighter transcriptional control of the sensor apparatus (ratio of maximum to basal GFP fluorescence around 3) compared to the Pseudomonads (ratio of maximum to basal GFP fluorescence around 1.5).

While the sensor portion of the device behaved similarly in each chassis, the memory apparatus – measured via RFP fluorescence – performed quite differently and showed significant chassis effect. Both *Pseudomonas* hosts performed significantly better than the *E. coli* counterparts (Fig. 2a). They had good transcriptional control and strong RFP fluorescence – *Pp* and *Pf* respectively 3- and 6-times more than the next best *E. coli* host, *EcN*. Interestingly, *Ec* was the lowest performing host from this study, both in terms of dynamic range and maximum fluorescence. In fact, it had a maximum RFU about 24-times lower than the best performing host, *Pf*.

Another important performance metric for cross-chassis comparison was the time scale of induction (Fig. 2b). This was quantified by the activation coefficient, which is the time for fluorescence to reach half maximum at a given IPTG concentration (21). These had very different profiles than maximum relative fluorescence measurements. *EcN* showed the fastest induction time, but was largely flat with respect to IPTG concentration. This effect was quantified by comparing half saturation times at 0.01 mM IPTG (6.17 ± 0.17 h) and 1 mM IPTG (6.26 ± 0.32 h). For others, the activation coefficient decreased with increasing IPTG concentration and approached a minimum value at the highest IPTG concentrations. Minimum GFP half saturation times were 11 ± 0.17 h, 12.34 ± 0.096 h and 15.09 ± 0.22 h for *Pf, Ec* and *Pp* respectively at 1 mM IPTG. RFP fluorescence followed a similar pattern, corresponding directly to induction of *P*_*lac*_ by IPTG. This result is evidence that the expression of Bxb1 – thus *P*_*lac*_ strength – was the rate limiting step in the process from induction by IPTG to DNA flipping.

### Population-based comparisons

Population-level measurements of the device was also carried out by flow-cytometry at five time points during the growth of the microbes (Fig. 3). All populations showed a portion with slight RFP fluorescence (slight tailing of the peaks) from the initial point of induction until about 4-6 hr time period. However, at low IPTG concentrations (0 and 0.01 mM), the RFP populations decreased in most of the species (observed by a sharpening of the peaks) until about 8-16 h. This observed effect evinced an initial and small population harboring a flipped memory element, that were quickly overtaken by the unflipped population because of low level of induction. At later time points and at higher IPTG concentrations, the populations shifted towards more flipped state.

**Figure 3.**
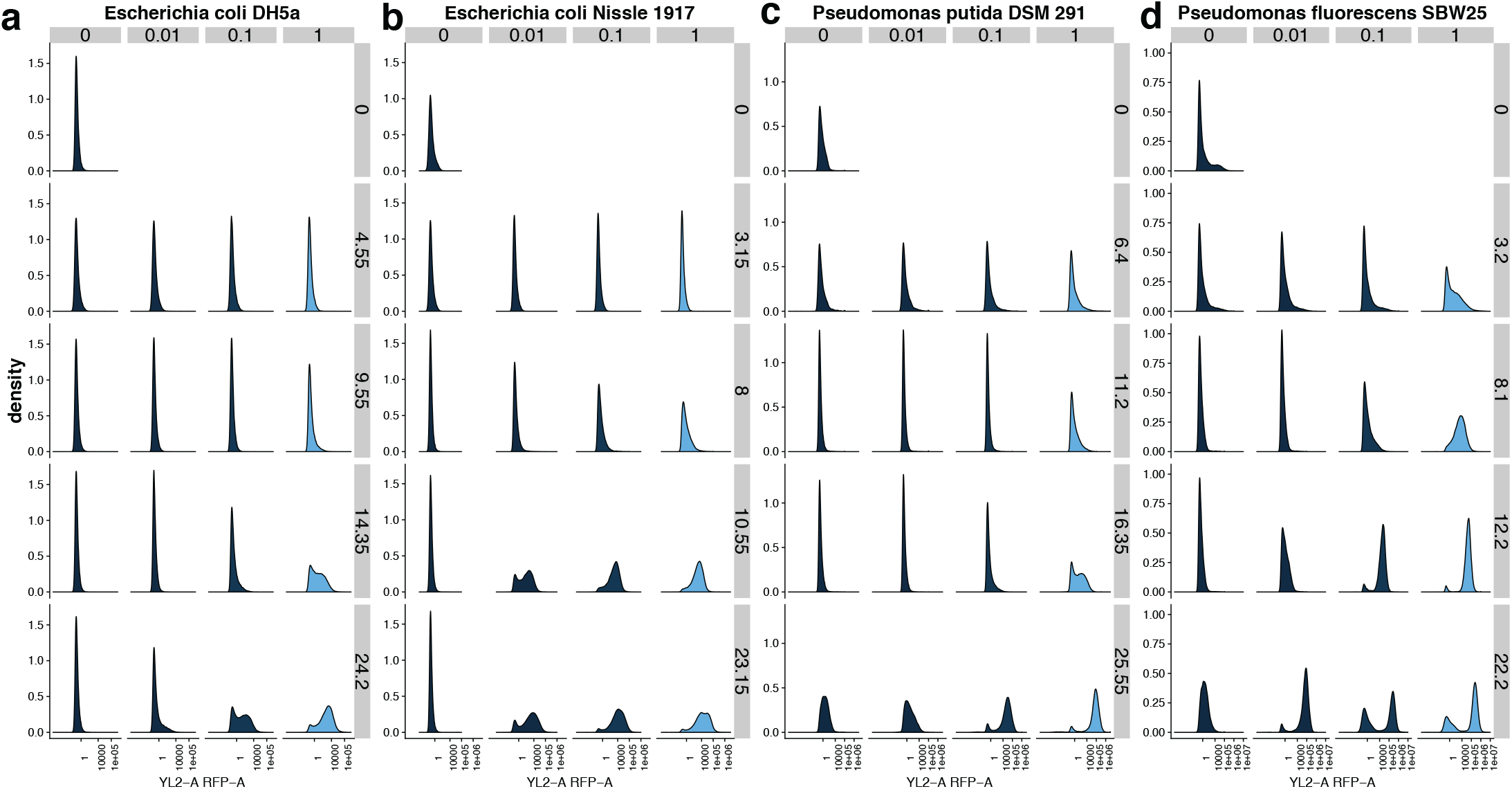
Population-based measurements of RFP reporting of signal recording across hosts. (a) Escherichia coli DH5⃇, (b) Escherichia coli Nissle 1917, (c) Pseudomonas putida DSM 291 and (d) Pseudomonas fluorescence SBW25. The rows in each of the panels correspond with sampling times in hours and columns are the IPTG concentrations given in mM. The first row in each plot shows data from the induction state from each experiment (0 h).

These measurements also show that transcriptional control and/or stability of the device was better in the *E. coli* hosts. The 0 mM IPTG treatments showed that the population distributions remained essentially constant for *Ec* and *EcN*, but both *Pp* and *Pf* shifted significantly at the final time point to favor an increasing number of cells with flipped memory element. This represents a false trigger of the event logger after extended periods (likely during stationary phase of growth) and is consistent with the interpretation of leaky expression of the *P*_*lac*_ promoter (Fig. 2a).

### Simulations and performance metrics across chassis

A kinetic model was formulated to help quantify how individual components of the event logger performed across each chassis. Like the physical construction of the device the model was broken down into the respective sensor and memory component categories. The sensor part of model included expressions that accounted for IPTG induction of the Bxb1 integrase (given as *I*) and GFP as shown by Equations 1 and 2.

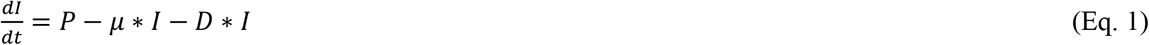

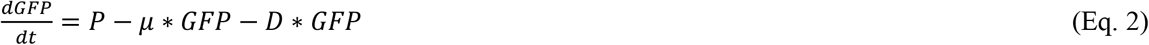

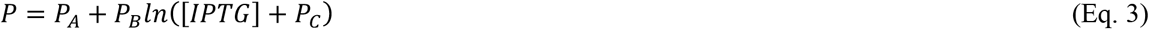

The relationship between IPTG concentration and promoter activity is given by Equation 3, where *P* represents the activity of *P*_*lac*_. It was found that this type of behavior adequately described the fluorescence output from the device. *D* is the protein degradation constant and *µ* is the specific growth rate, which accounts for dilution effects incurred by cell growth.

The memory component of the device was modeled by Equations 4 and 5, where *PB* is the fraction of un-flipped DNA and *LR* is the fraction of flipped DNA; *k*_*flip*_ is the rate constant for integrase-mediated recombination (flipping). We assumed that the plasmid copy number of each host was equivalent and that there existed an un-flipped induction state at the beginning of the experiment.

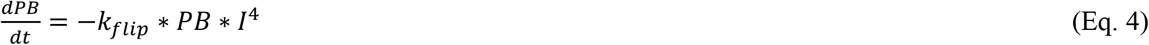

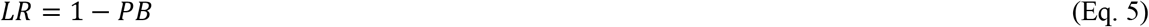

The rate constant, *k*_*flip*_ encompasses three-time steps: 1) the time required for the integrase (*I*) to form a tetramer; 2) the time required for binding of the tetramer to the DNA (*PB*) and 3) the DNA flipping event. The overall readout from the memory component of the device was given by expression of RFP (Equation 6), which is analogous to Eq. 2 describing GFP, with the exception of the non-inducible *tac* promoter (*P*_*RFP*_).

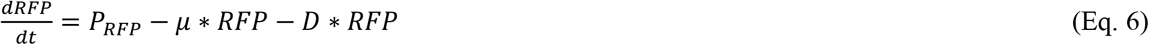

This model adequately explained the operation of the genetic device and fitted distinct parameters for each respective host. This was especially evident by the degree to which the model could be fit to the GFP and RFP time series data; each respective output from the sensor and memory components of the device (Fig. 4ab). However, the model’s ability to capture the dynamics of DNA flipping was variable between each of the hosts (Fig. 4c). We observed significant scatter derived from the qPCR assays that were designed to measure the orientation of the barcoded DNA associated with the digital memory read-out. These data were collected to determine the fraction of flipped DNA. The noise in the measurement likely resulted from plasmid degradation, variability of plasmid recovery and purification from each host. Yet, the model still conveyed the overall the pattern for which IPTG induction instigated barcode flipping for each of the hosts and did a reasonably good job at fitting most of the data derived from each time series measurement.

**Figure 4.**
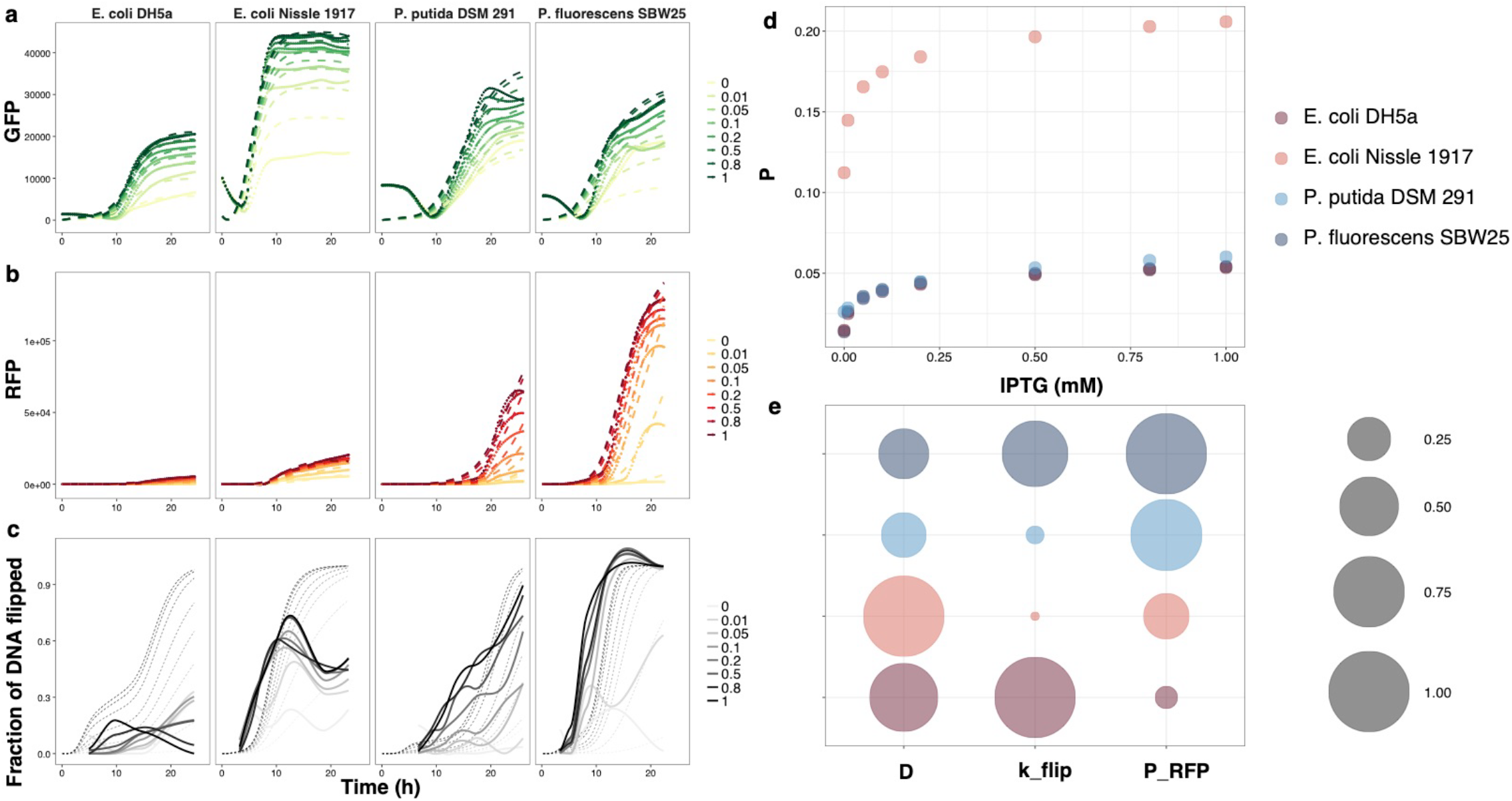
Comparative kinetics show chassis-dependent performance. The model outputs (dotted lines) of (a) the sensor component as fit by GFP fluorescence data; (b) the memory component as fit by RFP fluorescence; and (c) the memory component as fit by the fraction of DNA flipped (qPCR measurements). Each simulated time trace (panels a through c) are overlaid on the respective data typed used to parameterize the model (solid lines). (d) The promoter strength of Plac (given in the model as P) calculated from estimated model parameters, PA, PB and PC, plotted against IPTG concentration. (e) Comparison of simulated kinetic parameters for the protein degradation constant (D); flipping constant (kflip) and strength of the constitutive Ptac promoter (PRFP) compared across each host.

Overall, the model helped show that the *Pseudomonas* hosts had favorable kinetics for operating this device despite the fact the genetic parts have been largely developed and optimized in *E. coli*. They had similar GFP promoter strength (P) as *Ec* (Fig. 4d) but much higher RFP promoter strength (*P*_*RFP*_) and lower degradation constant (*D*) than both the *E. coli* species (Fig. 4e). However, the rate of DNA flipping, *k*_*flip*_, was quite low in *Pp* which indicated a slightly reduced performance in this host. To our surprise, the simulations suggest that *Ec* – the benchmark chassis – had the most unfavorable kinetics (low *P*, *P*_*RFP*_ and high *D*). The probiotic strain, *EcN* showed the strongest ability to operate the sensor component of the device and was unique with respect to the suite of hosts tested in this study (Fig. 4d). This could be ascertained by combining the modeled predictions with experimental measurements. For instance, the simulated promoter strength of *P*_*lac*_ (see Eq. 3) showed that *Ec, Pp* and *Pf* are aligned with similar profiles, while the values of *P* for *EcN* were estimated to be about 4-times higher for any given concentration of IPTG. This result was consistent with independent and direct measurements of specific growth rates (Supplemental Fig. S1); *EcN* showed the fastest specific growth rate and should therefore have the highest dilution of expressed GFP protein leading to decreased fluorescence. Yet, *EcN* also showed the highest GFP fluorescence. Thus, the strength of *P*_*lac*_ would need to be much higher to account for these opposing effects – as found by the model.

## DISCUSSION

Synthetic biologists commonly (re-)discover that even the most well-characterized genetic parts often will not function in a predictable manner when taken out of the context from which they were originally characterized. For any given host, this unpredictability can arise from interference between genetic parts that have been introduced as well as cellular noise inherent to the native biological system (22,23). Yet, the degree to which these factors are influenced by the biology of any given microbial species requires that the same genetic parts be used and compared across multiple hosts. Here, we showed that an identical genetic device can be constructed in a broad-host-range vector and ported across multiple microbial species and that its performance is host-dependent.

We chose to deploy a relatively simple event logger and experimental design, which has enabled this study to demonstrate emerging capability of broad-host-range genetic devices. In fact, the Bxb1 serine integrase was originally harnessed by synthetic biologists (24) in part because it does not require host cofactors, which is a feature enabling reuse across multiple hosts (25,26). The results from this current study confirm the suitability of Bxb1 as a broad-host-amenable genetic part by comparatively quantifying its performance with our broad-host event detector expressed from four species. While it is clear that more species and devices need to be tested before more extensive broad-host-range parts libraries can mature, this early step is an important contribution towards alleviating our current dependency and limitations on small subset of model microbial hosts.

Despite the fact that *E. coli* DH5α is a common tool for the design and implementation of modern genetic devices, we found that in many ways it was the least ideal host for the implementation of a device for actual application as tested in this study. In fact, a primary finding was that – compared to *Ec –* the two *Pseudomonas* species (*Pp* and *Pf*) showed reasonable potential as chassis for chemical event logging even though the majority of previously published reports on the parts used to build the device have only considered *E. coli* (3,4,6,27). We regard this as a promising result because these and closely related *Pseudomonads* are known to have tremendous metabolic potential for the synthesis of novel compounds (28,29), consuming complex substrates (30,31) and persisting in a wide range of habitats that include soils, plant tissues and marine ecosystems (32–35). Expanding the synthetic parts list for *Pseudomonas* species will undoubtedly enable new biotechnological applications that should include chemical sensing and event logging in complex natural environments.

*E. coli* Nissle 1917 was also chosen for comparative analysis in this study because it served as an intra-species comparator to *Ec*. It was also chosen because of its growing importance in the bio-design community based on the fact that it is a commonly used probiotic(36) and highly genetically tractable. Researchers are rapidly uncovering many exciting opportunities to use *EcN* and other probiotic-hosts as programable therapeutic agents and/or diagnostic tools for human health(37–39). In some cases, differences in intra-species performance – within *E. coli* strains – exceeded inter-species variability. This was somewhat unexpected and specifically evident from comparisons made on the sensor component of the device, which performed better in *EcN* as compared to *Ec* and both *Pseudomonas* hosts. This was specifically evident by comparing the kinetics associated with the sensor apparatus and indicates that *EcN* maintained the tightest control and largest dynamic ranges of the IPTG inducible components of the device.

Kinetic parameters estimated from the model provided a good quantitative comparison of biological properties that cannot be easily measured (Fig. 4e). For instance, the degradation constant, *D*, is found to be fairly similar in all the hosts, which is hardly surprising considering the standardized growth conditions and similar growth rates. Estimates of the flipping rate constant (*k*_*flip*_), however, were highly variable. In contrast to *D*, which is more indicative of cellular physiology, *k*_*flip*_ is more representative of the device-specific kinetics. This parameter depends on a number of biological factors such as codon usage, transcription, translation, protein folding as well as the efficiency of Bxb1-mediated recombination. Based on the model-enabled predictions, *EcN* stood out with a very small *k*_*flip*_ values; about 89 times smaller than the largest value attributed to *Pf*. Its high transcription rate (given by estimates of *P*) and low *k*_*flip*_ account for the observation of fast DNA flipping after initial induction followed by relatively immediate saturation (Fig. 4c). The *k*_*flip*_ values of Ec and *Pp* are moderate but the reason for the higher value of *Ec* relative to *Pp* is still somewhat uncertain since the fraction of DNA flipped is higher in *Pp* than *Ec*. The fourth parameter, *P*_*RFP*_, is the measure of the strength of *P*_*tac*_ promoter and varies in the same way as RFP fluorescence. This promoter was actually found to work better – as assayed by the strength or RFP fluorescence – in the *Pseudomonads* than *E. coli* species.

Integrated data and kinetic modelling approaches are useful for quantifying and comparing performance across hosts. One limitation, however, was that our approach contained few species-specific physiological parameters. The exception to this is the specific growth rate (μ). Although the hosts in this study all showed similar growth rates, the specific growth rate should prove to be an important consideration when evaluating the performance of a device as hosts and growth conditions change. It was also interesting that we were able to observe and simulate dynamics in the device’s performance while the cells were in stationary phase. Often, experimental observations made on engineered devices are only contextualized during log growth phases. However, future applications such as chemical event logging in dynamic environments will be better served by understanding how chassis/device pairs may function through lag, log and stationary phases of growth. This is a point that shall require more deserving attention in future studies.

The field of microbial bio-design is keen to harness new, non-traditional hosts for synthetic biology applications. Some significant advancements towards programing genetic devices – including sensors – have already been shown in other non-traditional microbial hosts. Of specific note are previous success shown in a human gut microbe *Bacteroides thetaiotaomicron* (40) and a suite of proteobacteria isolated from a bee gut microbiome (41). Here in this current study, we have advanced an emerging concept of broad-host-range genetic devices. While this is certainly a new frontier, some notable examples have preceded this current report including a study by Kushwana and Salis that that presented the concept of “portable power supplies” between species and demonstrated that some genetic parts can ported between *E. coli*, *P. putida* and *Bacillus subtilis* (14). Another important avenue has been the pursuit of broad-spectrum genetic parts such as the promoters presented in a study from Yang et al. that are operational between *E. coli*, *B. subtilis* and *S. cerevisiae* (15). The efforts to date – including our current study – have only considered a relatively small set of microbes. Future developments on cross-chassis devices may encounter new technical hurdles as the taxonomic diversity of hosts are expanded. Once harnessed, the concept of broad-host-range genetic devices should also bring new species-specific-applications. The major technical hurdle that will need to be overcome for developing chemical sensing capabilities will be the discovery or engineering of genetic components with specificity for analytes of real-world interest. The current suite of commonly used transcriptional factors and inducible promoters are clearly limited. New parts discovery and characterization efforts are sorely needed to advance the current state of microbial biodesign.

### Conclusions

We quantified the chassis effect of an integrase-based chemical event logger across multiple species and two different Genera – *Pseudomonas* and *Escherichia*. The performance of sensor and memory components changed according to each host as ascertained via integrated experimental measurements and outputs from kinetic models. Specifically, *EcN* – a common probiotic bacterium – showed the tightest control and most stability of the sensor apparatus that regulated expression of the Bxb1 integrase. Both *Pseudomonas* hosts showed greater RFP output signals that corresponded with Bxb1-mediated recombination in the memory component of the device. A primary finding of this study was that – compared to *Ec –* the two *Pseudomonas* species (*Pp* and *Pf*) showed reasonable potential as chemical event logging chassis. This study advances an emerging frontier in synthetic biology that aims to build broad-host-range devices and understand the context by which different microbial species can execute programmable genetic operations

## SUPPLEMENTARY DATA

The raw data for this study along with and the sequence information for the genetic construct pB2lacBxb1G-R (shown in Fig. 1b) are available from the Open Science Framework (OSF) under the name “A broad-host-range event detector: data and models” (DOI 10.17605/OSF.IO/J295C) at https://osf.io/j295c/. This OSF project also contains the R Markdown scripts that can be used to reproduce all of the analyses, statistics and technical graphs presented in this manuscript.

## ACKNOWLEDGMENTS

This research was supported by funding from the National Security Directorate Seed Initiative, a Laboratory Directed Research and Development (LDRD) Program of Pacific Northwest National Laboratory (PNNL). PNNL is operated for the DOE by Battelle under contract no. DE-AC05-76RLO-1830. Specific acknowledgments are given to Rose Perry at PNNL and John Repass at Genetics for technical assistance with graphics and real time PCR. The authors would also like to thank Drs. Victoria Hsiao and Richard Murray for kindly sharing materials and knowledge.

## Supplemental Information

**Table 1.**
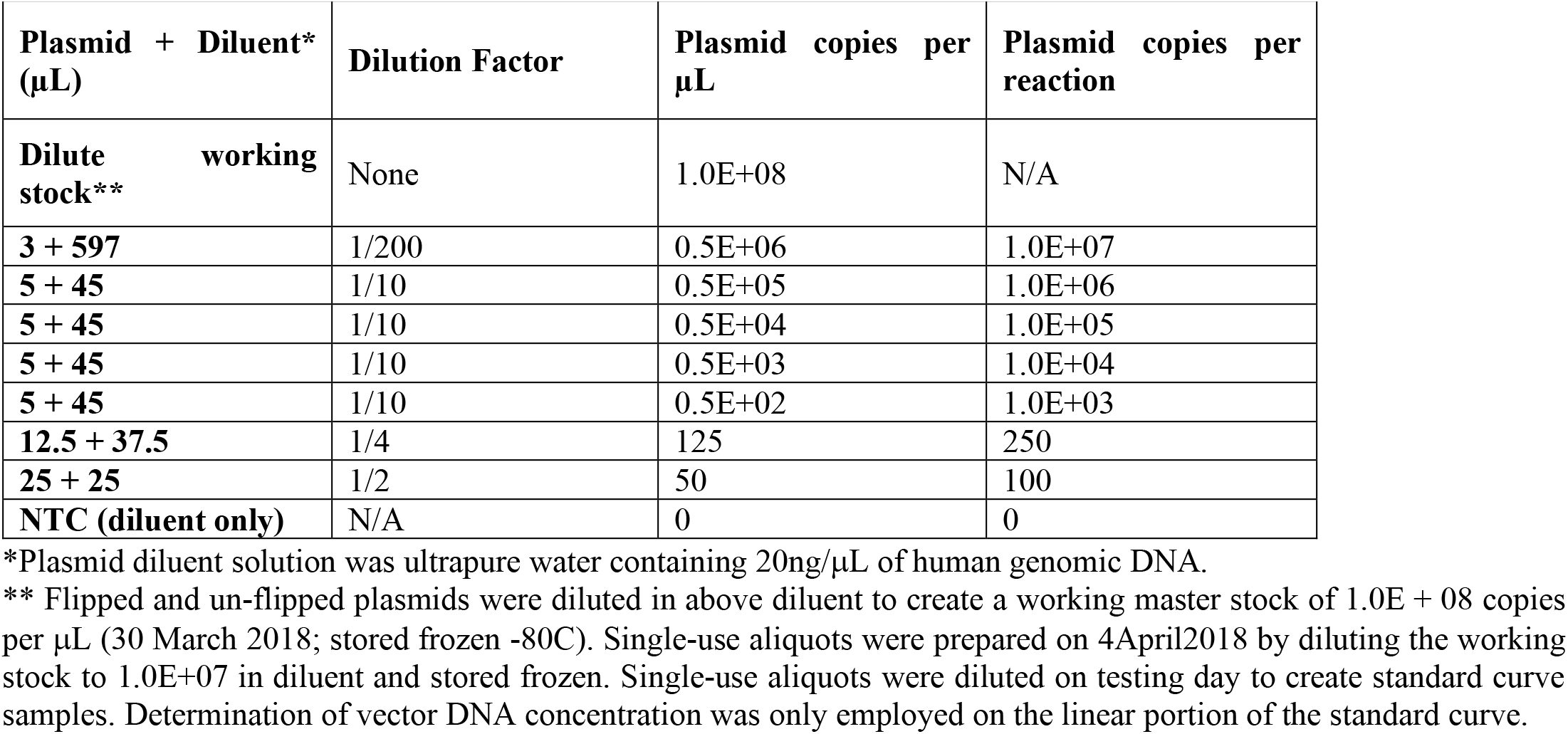
Standard Curves for qPCR -preparation of plasmid DNA serial dilutions for standard curve reactions

**Table 2.**
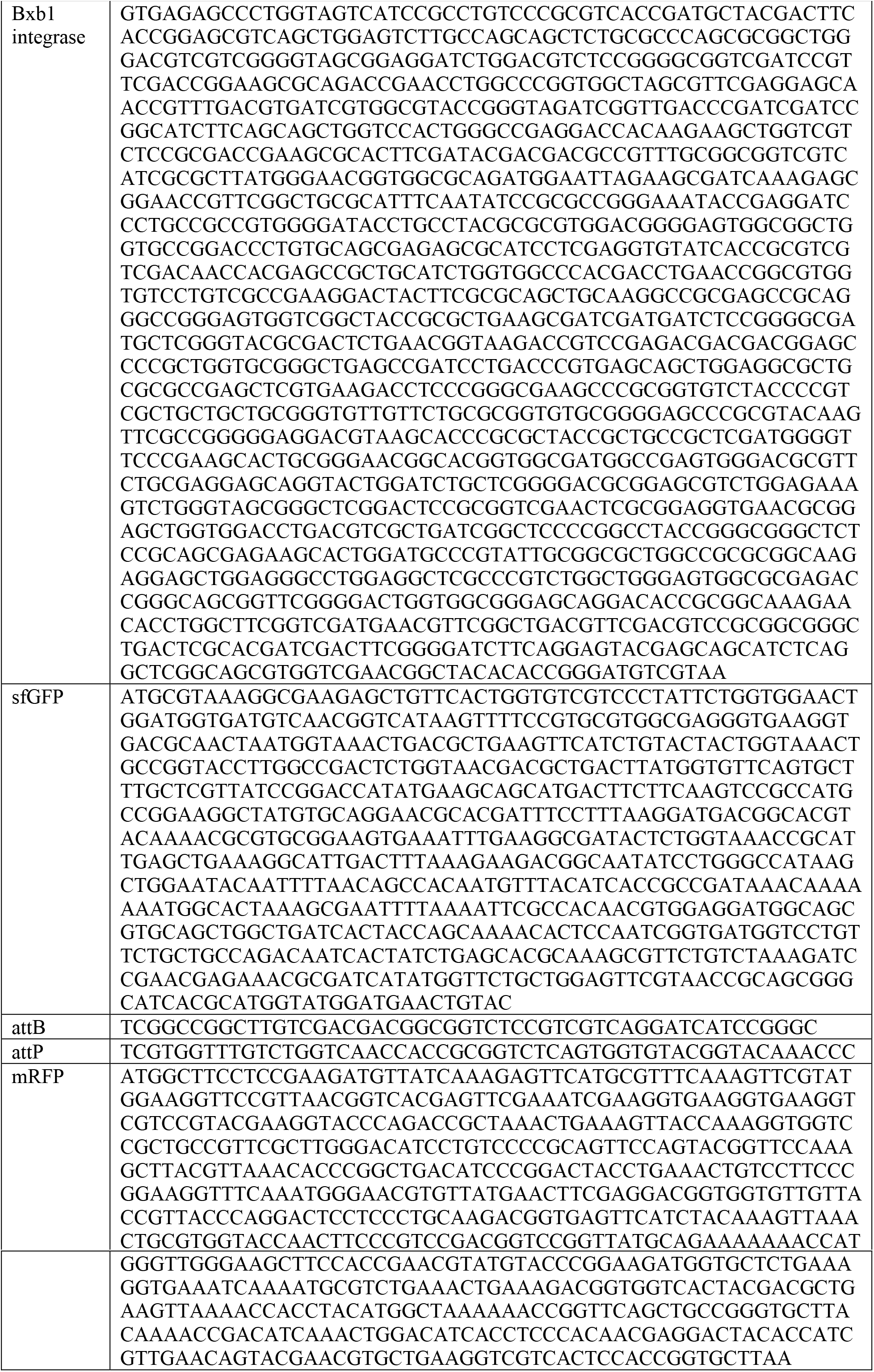
Sequences of major components

**Table 3.**
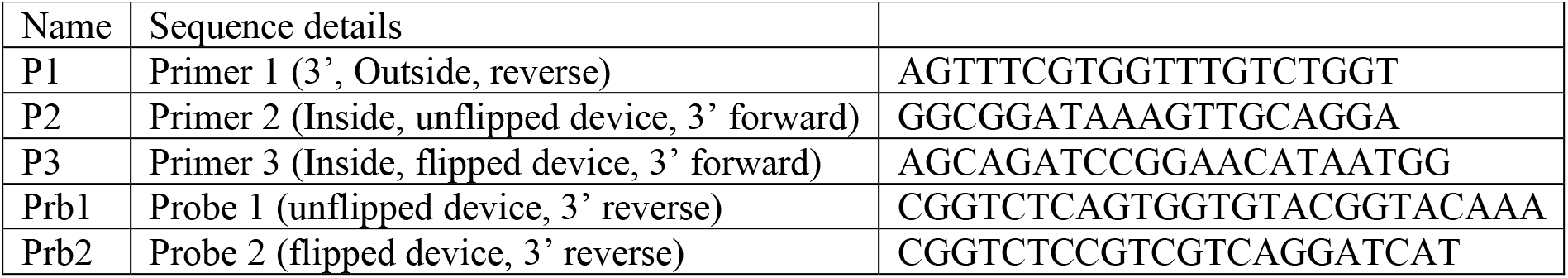
qPCR primer/probe sequences

**Supplemental Figure S1.**
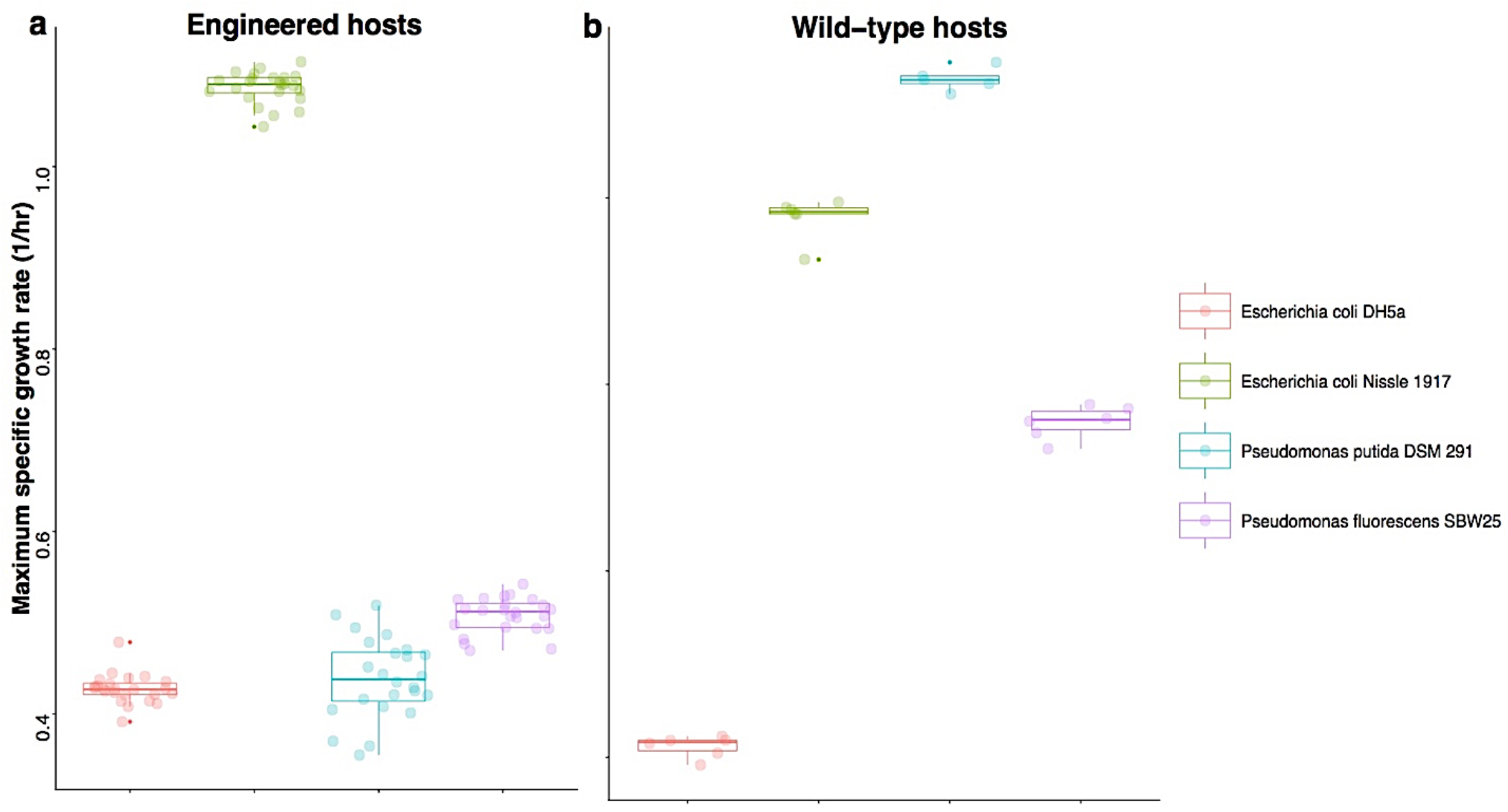
Comparisons of the specific growth rates between hosts. **(a)** The maximum specific growth rates of each host expressing the pBBR1MCS2-based event logging device at all IPTG concentrations tested. **(b)** Maximum specific growth rates of each wild-type host; absent of expression vector and device.

**Figure S2.**
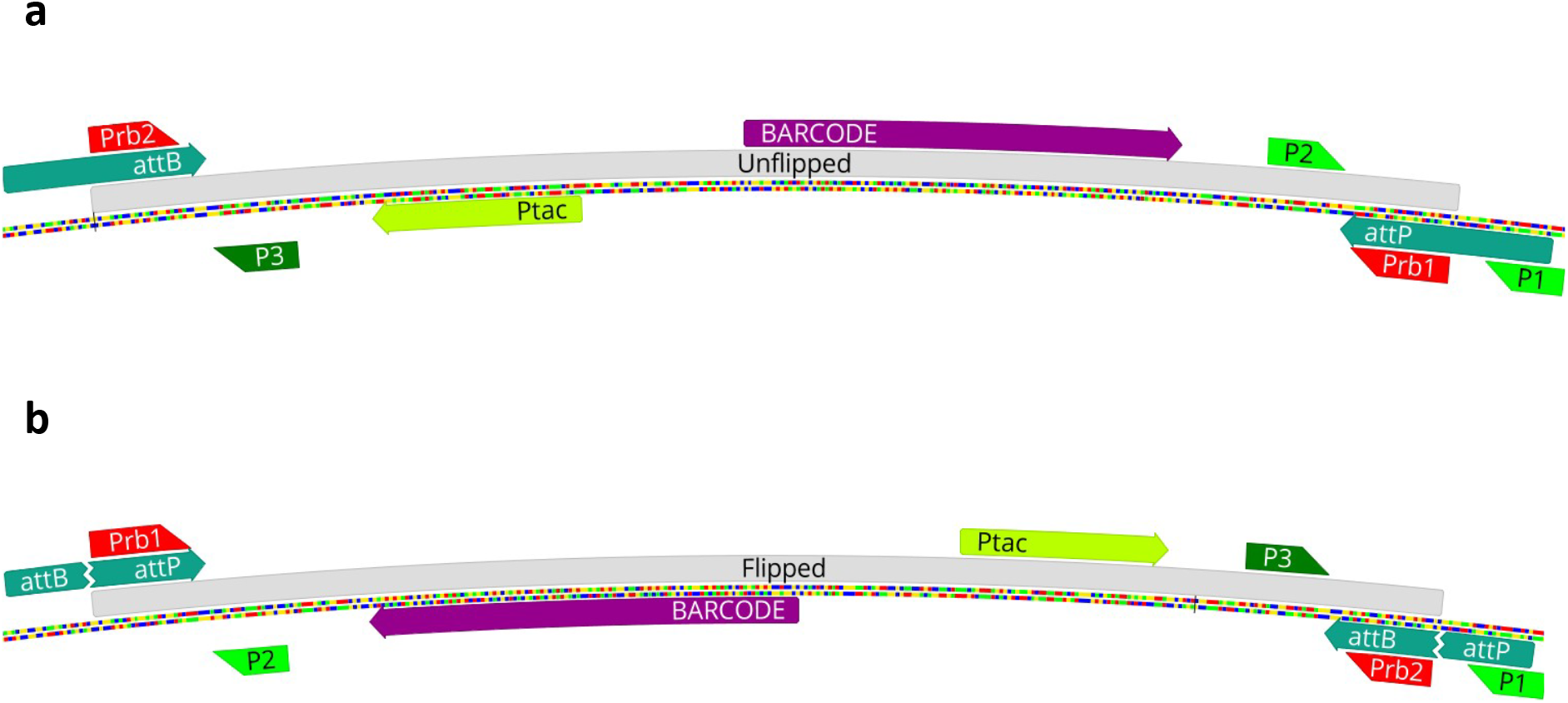
Device maps with primer, probe locations - (a) unflipped, (b) flipped. In the unflipped orientation primers P1 and P2 form a pair and amplifies, while P3 remains unpaired and causes very little amplification. When the device is flipped, positions of P2 and P3 are reversed, and P3 now forms a pair with P1 and amplifies.

